# Revealing Functional Hotspots: Temperature-Dependent Crystallography of K-RAS Highlights Allosteric and Druggable Sites

**DOI:** 10.1101/2025.02.27.639303

**Authors:** Samuel L. Deck, Megan Xu, Matt Stankus, Shawn K. Milano, Richard A. Cerione

## Abstract

K-RAS mutations drive oncogenesis in multiple cancers, yet the lack of druggable sites has long hindered therapeutic development. Here, we use multi-temperature X-ray crystallography (MT-XRC) to capture functionally relevant K-RAS conformations across a temperature gradient, spanning cryogenic to physiological and even “fever” conditions, and show how cryogenic conditions may obscure key dynamic states as targets for new drug development. This approach revealed a temperature-dependent conformational landscape of K-RAS, shedding light on the dynamic nature of key regions. We identified significant conformational changes occurring at critical sites, including known allosteric and drug-binding pockets, which were hidden under cryogenic conditions but later discovered to be critically important for drug-protein interactions and inhibitor design. These structural changes align with regions previously highlighted by large-scale mutational studies as functionally significant. However, our MT-XRC analysis provides precise structural snapshots, capturing the exact conformations of these potentially important allosteric sites in unprecedented detail. Our findings underscore the necessity of advancing tools like MT-XRC to visualize conformational transitions that may be important in signal propagation which are missed by standard cryogenic XRC and to address hard-to-drug targets through rational drug design. This approach not only provides unique structural insights into K-RAS signaling events and identifies new potential sites to target with drug candidates but also establishes a powerful framework for discovering therapeutic opportunities against other challenging drug targets.

## Introduction

Small GTPases are a class of guanine-nucleotide-binding proteins that act as molecular switches, regulating critical cellular processes such as growth, proliferation, differentiation, and migration in response to extracellular signals (1–3). These proteins transition dynamically between an inactive GDP-bound state and an active GTP-bound state, driven by GDP-GTP exchange and GTP hydrolysis. This activation cycle is tightly regulated by guanine nucleotide exchange factors (GEFs), which facilitate GDP release and GTP binding, and GTPase-activating proteins (GAPs), which catalyze the hydrolysis of GTP back to GDP (4,5). The structural changes induced during this cycle are crucial for downstream signaling and the coordination of vital cellular activities (6).

Among the small GTPases, K-RAS (Kirsten rat sarcoma viral oncogene homolog) has garnered significant attention due to its pivotal role in human cancers (7). Mutations in K-RAS are observed in approximately one-third of all malignancies, with substitutions at key residues, such as glycine 12 (G12), leading to its constitutive activation (8,9). This leads to a persistent activation of KRAS’ downstream signaling pathways which drives oncogenesis and makes K-RAS a high-priority therapeutic target (10). However, K-RAS has long been deemed “undruggable” due to its smooth surface and lack of well-defined binding pockets, posing substantial challenges for rational drug design (11). Despite the recent development of mutation-specific inhibitors, such as covalent G12C-targeting compounds, there is still an urgent need for innovative strategies to uncover additional druggable sites and expand therapeutic options (12–14).

Cryogenic crystallography has been the standard tool for elucidating the structural details of small GTPases, including K-RAS, and has provided important insights into their function and inhibitor interactions (15–18). However, this technique has notable limitations. The freezing process immobilizes proteins in static conformations, potentially obscuring biologically relevant structural dynamics and introducing artifacts that may skew interpretations (19–22). As K-RAS function relies on subtle conformational changes within flexible regions such as Switch I, Switch II, and the P-loop, these limitations could hinder the identification of key structural transitions as well as fail to detect critical druggable sites and impede drug development efforts.

To address these challenges, multi-temperature X-ray crystallography (MT-XRC) offers a promising solution by capturing protein structures across a range of temperatures, from cryogenic to physiological and even “fever” conditions (23–25). This approach enables the observation of temperature-dependent conformational dynamics, providing a more accurate representation of the protein’s behavior under physiological conditions (26–28). By revealing hidden structural features, such as transient conformational changes that yield new allosteric pockets, MT-XRC can uncover key regions for inhibitor binding that are not apparent in cryogenic structures (27,28). Previous studies from our laboratory and others have demonstrated the value of room-temperature crystallography in resolving critical conformational differences, offering new insights into protein function and drug binding (29).

In this study, we apply MT-XRC to investigate the structural dynamics of wild-type and mutant K-RAS proteins, focusing on the oncogenic G12C mutant as a paradigmatic example. By analyzing high-resolution structures across a temperature gradient, we reveal a temperature-dependent conformational landscape that sheds light on critical structural features, including strained GDP versus GTP binding, dynamic rearrangements within the classical conformationally-sensitive ‘Switch domains,’ and the adaptability of key allosteric sites. Our findings demonstrate that MT-XRC provides a unique advantage in visualizing dynamic and physiologically relevant conformations, simplifying the identification of allosteric sites, and directly informing inhibitor design. These insights not only shed light on potentially important structure-function features of K-RAS but also establish MT-XRC as a powerful tool for tackling other hard-to-drug targets, paving the way for more effective therapeutic strategies in cancer treatment.

## Results

### Temperature-Driven Structural Perturbations Expose Ordered Conformational Heterogeneity in K-RAS

To capture conformational states of K-RAS that may be obscured under cryogenic conditions, we determined a 1.4 Å room-temperature (RT, 293 K) crystal structure of GDP-bound wild-type (WT) K-RAS. The structure yielded a complete and well-resolved model of the G-domain (residues 1–169), including the P-loop, Switch I, Switch II, and the hypervariable region, regions that are often incompletely resolved in cryogenic datasets owing to their intrinsic flexibility (Supplementary Fig. 1A–D) (30).

The RT structure preserves the canonical K-RAS fold and a well-defined nucleotide-binding pocket, with GDP and its associated magnesium ion coordinated by a conserved network of hydrogen-bonding and hydrophobic interactions (Supplementary Fig.1 B,C) (10,11,32). Comparison with the corresponding cryogenic structure solved at 100 K (PDB ID: 4OBE) showed that, although the overall fold is maintained, localized conformational differences emerge in functionally important regions (Supplementary Fig. 1D) (33).

These differences are not broadly distributed, but instead cluster within discrete regulatory elements of the protein. Residue-level changes map to the P-loop, Switch I, the inter-switch region, positions within the allosteric lobe, and the hypervariable region (Fig.1 A,B). Structural overlays reveal temperature-dependent side-chain rearrangements at representative sites, including Asp30 and Glu31, the inter-switch segment spanning Arg41–Leu52, and more distal positions such as Cys118, Asp119, Leu120, Pro121, Lys147, Glu168, and Lys169 (Fig. 1C–F). Together, these observations indicate that increasing temperature redistributes conformational sampling into defined surface hotspots rather than inducing global structural change.

**Figure 1.**
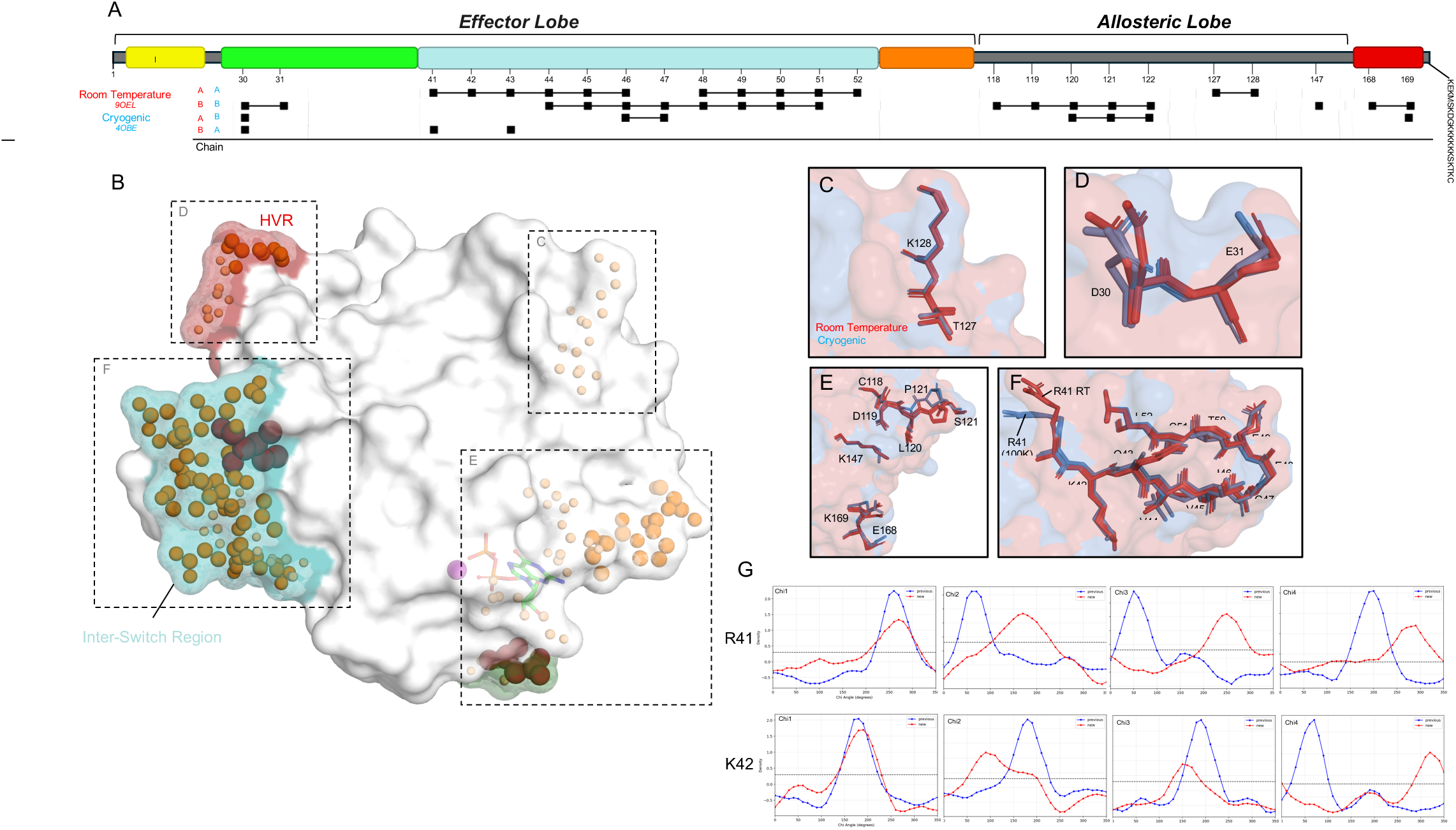
Comparison of room-temperature and cryogenic side-chain conformations in K-RAS. ***A,*** Schematic summary of residue-level conformational differences between the room-temperature and cryogenic WT K-RAS structures across the G-domain. Major structural elements are indicated, including the P-loop (yellow), Switch I (green), the inter-switch region (cyan), Switch II (orange), and the hypervariable region (HVR, red), grouped into the effector and allosteric lobes. Residues exhibiting temperature-dependent changes are mapped by position along the sequence. ***B,*** Surface representation of K-RAS highlighting clusters of residues with altered conformations between the two structures. Distinct regions of change are distributed across the inter-switch region, the HVR, and sites surrounding the nucleotide-binding surface; boxed areas correspond to the zoomed views shown in panels ***C–F***. ***C–F,*** Superpositions of representative residues and residue clusters from the room-temperature (red) and cryogenic (blue) structures, illustrating examples of temperature-dependent side-chain rearrangements. These views highlight local differences in rotameric state and side-chain placement within the surrounding protein surface. ***G,*** Side-chain dihedral angle plots for representative residues R41 and K42 comparing the room-temperature (red) and cryogenic (blue) structures across χ1–χ4, showing residue-specific shifts in rotamer preference between the two temperature conditions.

Several of these perturbations occur in regions central to nucleotide handling and effector engagement. The P-loop exhibits subtle outward displacement, consistent with increased conformational freedom at elevated temperature. Switch I and Switch II likewise display enhanced variability, and Arg41 adopts a more open conformation in the RT structure. More broadly, the inter-switch region undergoes coordinated rearrangement, consistent with a role as a dynamic hinge coupling regulatory surfaces within K-RAS. This redistribution is further supported by side-chain dihedral analysis, which shows distinct shifts in rotamer preference for Arg41 and Lys42 between the RT and cryogenic structures (Fig. 1G).

Consistent with these structural changes, comparison of the RT and cryogenic WT structures shows increased flexibility across major regulatory regions, including the P-loop and Switch segments (Supplementary Fig. 1D). These findings support the view that cryogenic data collection can suppress conformational heterogeneity and obscure substates accessible under more physiological conditions (19–22). Taken together, these results show that RT crystallography captures an expanded but ordered conformational ensemble in WT K-RAS, revealing localized structural heterogeneity in regions linked to nucleotide exchange, effector recognition, and allosteric communication (23–29).

### Temperature-Sensitive Conformational Shifts in K-RAS G12C

Building on our observations of temperature-dependent flexibility in WT K-RAS and the success in determining the structure at RT, we next examined the K-RAS G12C mutant, a clinically significant variant due to its prevalence in multiple cancers and its unique druggable properties (13). The G12C mutation introduces a reactive cysteine at position 12, altering the nucleotide-binding pocket and nearby regions in ways critical for both its oncogenic function and drug interactions. To assess the structural dynamics of the G12C mutant at RT, we compared it to the RT structure of WT K-RAS. Overall, the global fold remains conserved between the two structures (Fig. 2A). However, the G12C mutation in K-RAS induces structural perturbations that affect key residues involved in nucleotide binding and overall conformational stability. Structural comparisons between the WT and the G12C proteins at room temperature reveal distinct alterations at residues 1, 4, 23–26, 30, 46–51, 121, 122, 126, and 169. Notably, residues within Switch I (23–26) and Switch II (46–51) exhibit shifts in backbone positioning, which may influence their interactions with guanine nucleotides and effector proteins. Residues 1, 4, and 30 also display subtle deviations, suggesting potential alterations in nucleotide coordination. Additionally, minor rearrangements in the α3-helix (residues 121, 122, 126, and 169) suggest a redistribution of structural strain that could impact the dynamic behavior of the protein.

**Figure 2.**
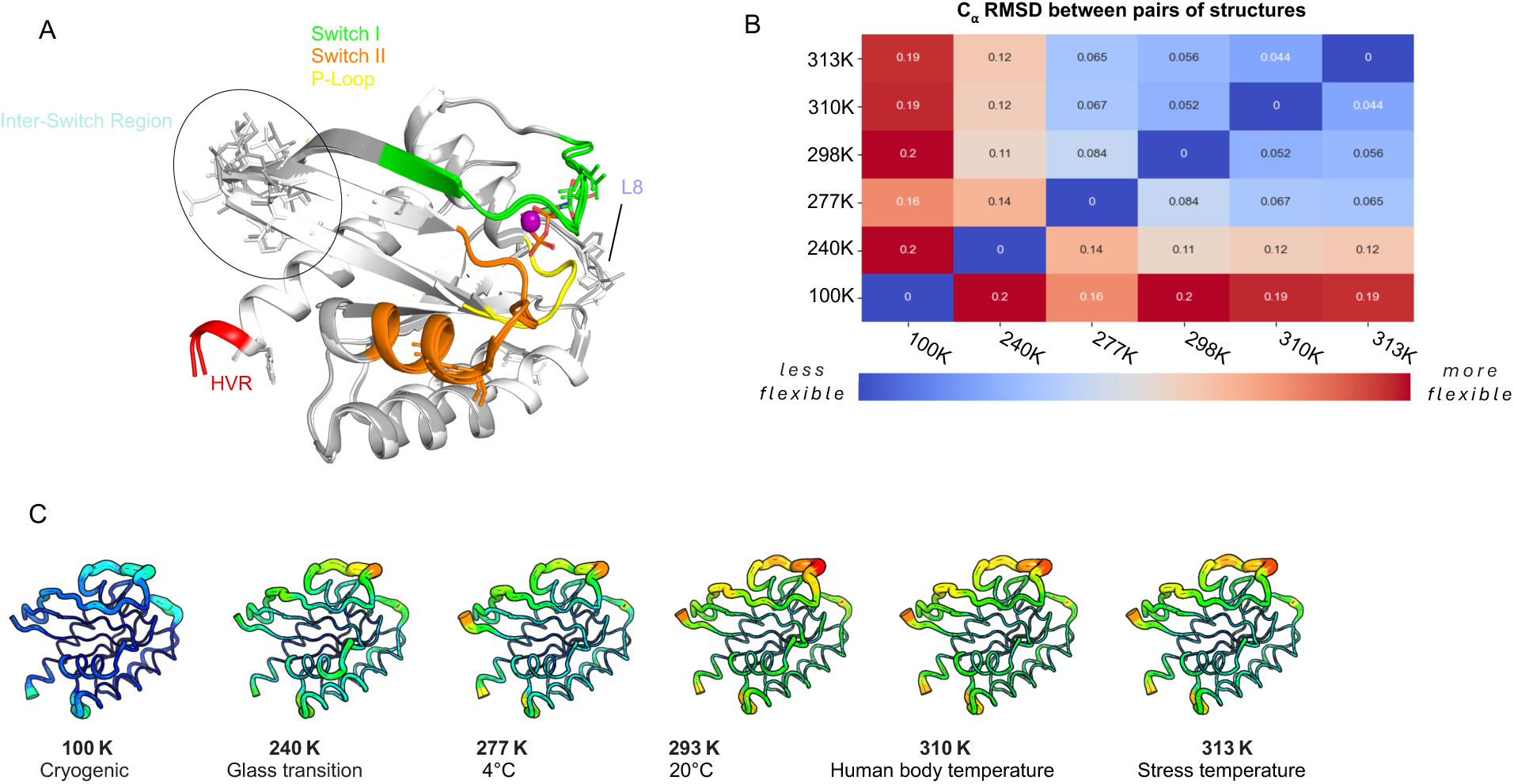
Multi-temperature Crystallography (MT-XRC) of the K-RAS G12C Mutant. ***A***, Overlay of the WT K-RAS (white) and K-RAS G12C mutant (grey) structures. Switch I (green), Switch II (orange), P-loop (yellow), (HVR) (red), Inter-Switch (blue), and Loop 8 (dark blue) regions are highlighted. Structural residues differences in Inter-Switch region are shown and circled. ***B***, Table displaying the RMSD (Root Mean Square Deviation) for the G12C mutant K-RAS structures solved at various temperatures (100K, 240K, 277K, 293K, 310K, and 313K). Blue represents the lowest and red the highest B-factor flexibility. ***C***, Structures of the G12C K-RAS proteins are drawn in cartoon putty with rainbow coloring with Blue representing the lowest and red the highest B-factor values. The size of the tube also reflects the B-factor, with larger tubes indicating higher B-factors.

These various changes illustrate how the G12C substitution perturbs key regulatory elements within K-RAS, with structural deviations propagating to regions critical for nucleotide exchange and effector binding. To further explore temperature dependent conformational changes of this important K-RAS oncogenic mutant, we determined its high-resolution crystal structures across a temperature gradient, ranging from cryogenic (100K, −173°C) to physiological (310K, 37°C) and fever-like (313K, 40°C) conditions. A comparison chart (Fig. 2B) summarizes the root-mean-square deviation (RMSD) values for K-RAS G12C structures over these temperature ranges. This shows a clear increase in structural variability as the temperature increases. At 100K, the RMSD values are relatively low, indicating minimal structural change and reduced flexibility. As temperatures increase, the RMSD values for the G12C structures grow, showing a more pronounced increase in RMSD at higher temperatures. At 313K, G12C structures exhibit the highest RMSD values, reflecting greater conformational shifts in the G12C mutant compared to the WT protein. This is visually illustrated in a cartoon putty representation of the G12C mutant at different temperatures (Fig. 2C), in which we see that the P-loop, Switch I, and Switch II regions exhibit increased flexibility at higher temperatures. The width and color intensity of the putty representation correspond to the B-factor values, with thicker and more intensely colored regions indicating increased atomic motion and flexibility. At cryogenic temperatures, these regions appear relatively rigid. As the temperature increases, the P-loop, Switch I, and Switch II regions become more prominent indicating dynamic structural mobility. The enhanced flexibility in these regions suggests that the increased temperature induces conformational changes in K-RAS that can be important for interactions with downstream effectors and potential therapeutic drugs. These structural changes can also have significant consequences for drug design, as they highlight regions that may be more accessible or altered at physiological temperatures, thus offering new opportunities for targeting K-RAS G12C with specific inhibitors.

### Temperature-Dependent Pocket Formation in K-RAS G12C Structures

Given the temperature dependent structural perturbations induced by the G12C mutation, we sought to determine whether the structures for this oncogenic mutant over the temperature landscape would reveal novel binding pockets that potentially could be exploited for therapeutic intervention. Using FPocketWeb (34), we found that the RT structure had a total of 11 pockets, exceeding the number observed at both colder and warmer temperatures (Fig. 3C). This data suggests that temperature plays a critical role in the conformational plasticity of the G12C mutant. At cryogenic temperatures, seven pockets were detected, primarily aligning with known allosteric sites observed in previous studies (35). However, at physiological temperature, an additional eighth pocket emerged, corresponding to the groove that led to the development of the first covalent K-RAS G12C inhibitors (13). An overlay of our physiological-temperature structure for K-RAS G12C with the published structure for the K-RAS G12C-inhibitor complex (PDB: 4LUC) at this site shows that the pocket observed at 310K, 37°C closely matches the groove occupied by early covalent inhibitors (under stress conditions at 313K, 37°C this groove disappears) (Fig. 3B) (13).

**Figure 3.**
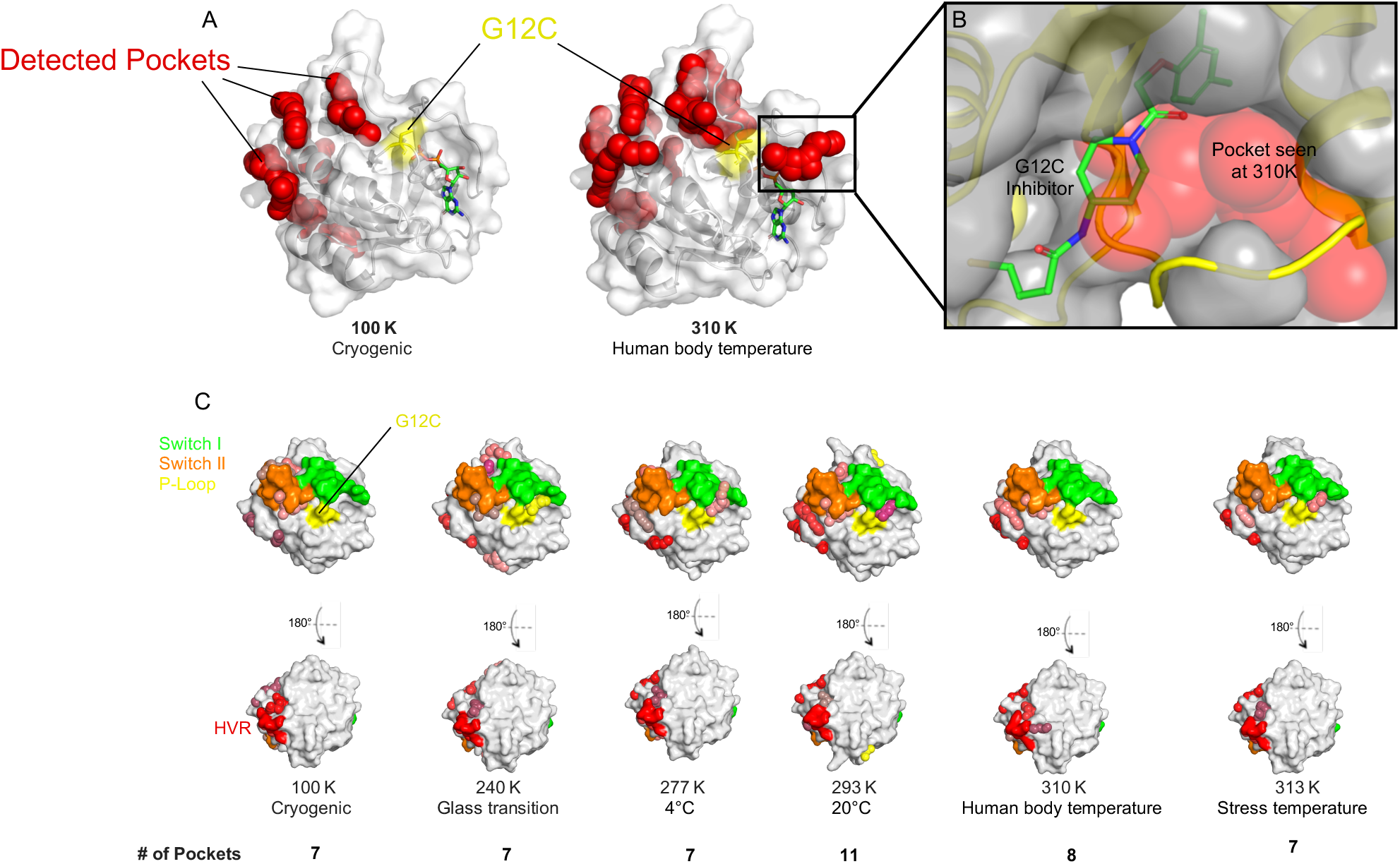
Temperature-dependent pocket formation in the K-RAS G12C structures. ***A***, K-RAS G12C structures at cryogenic (100 K) and physiological (310 K) temperatures are presented with detected pockets shown using FPocketWeb (red spheres), with residue 12 being highlighted in yellow. ***B***, Overlay of the cryogenic K-RAS G12C-inhibitor complex (yellow, cartoon) (PDB: 4LUC) with the 310K K-RAS G12C structure (grey, surface). The pocket that appears at 310K (red spheres) is the groove the inhibitor exploits to bind. ***C***, Surface representations of K-RAS G12C at various temperatures, with Switch I (green), Switch II (orange), P-loop (yellow), (HVR) (red), Inter-Switch (blue), and Loop 8 (dark blue) regions highlighted. Unique pockets are shown as colored spheres.

The presence of additional binding pockets at RT raises intriguing possibilities for targeting alternative transient allosteric sites that may not be apparent under more static conditions. These findings underscore the critical role of temperature in structural studies of K-RAS, suggesting that an optimal temperature during data collection may offer a more accurate representation of potential drug-binding sites and facilitate the design of next-generation inhibitors targeting oncogenic K-RAS.

### Conformational Analysis and Key Residue Flexibility Across Temperature Gradients

Given the importance of temperature-dependent structural dynamics in uncovering cryptic binding sites, we investigated whether specific regions of K-RAS G12C exhibit unique flexibility patterns that could indicate alternative allosteric or druggable sites. To quantify these temperature-driven conformational shifts, we analyzed the root mean square fluctuation (RMSF) of K-RAS G12C bound to GDP across different temperatures, plotting per-residue flexibility to identify regions of enhanced atomic displacement (36). Structural regions of interest are highlighted in shaded grey areas, indicating potential transient pockets or alternative drug-binding surfaces that may be occluded under cryogenic conditions (Fig. 4A).

**Figure 4.**
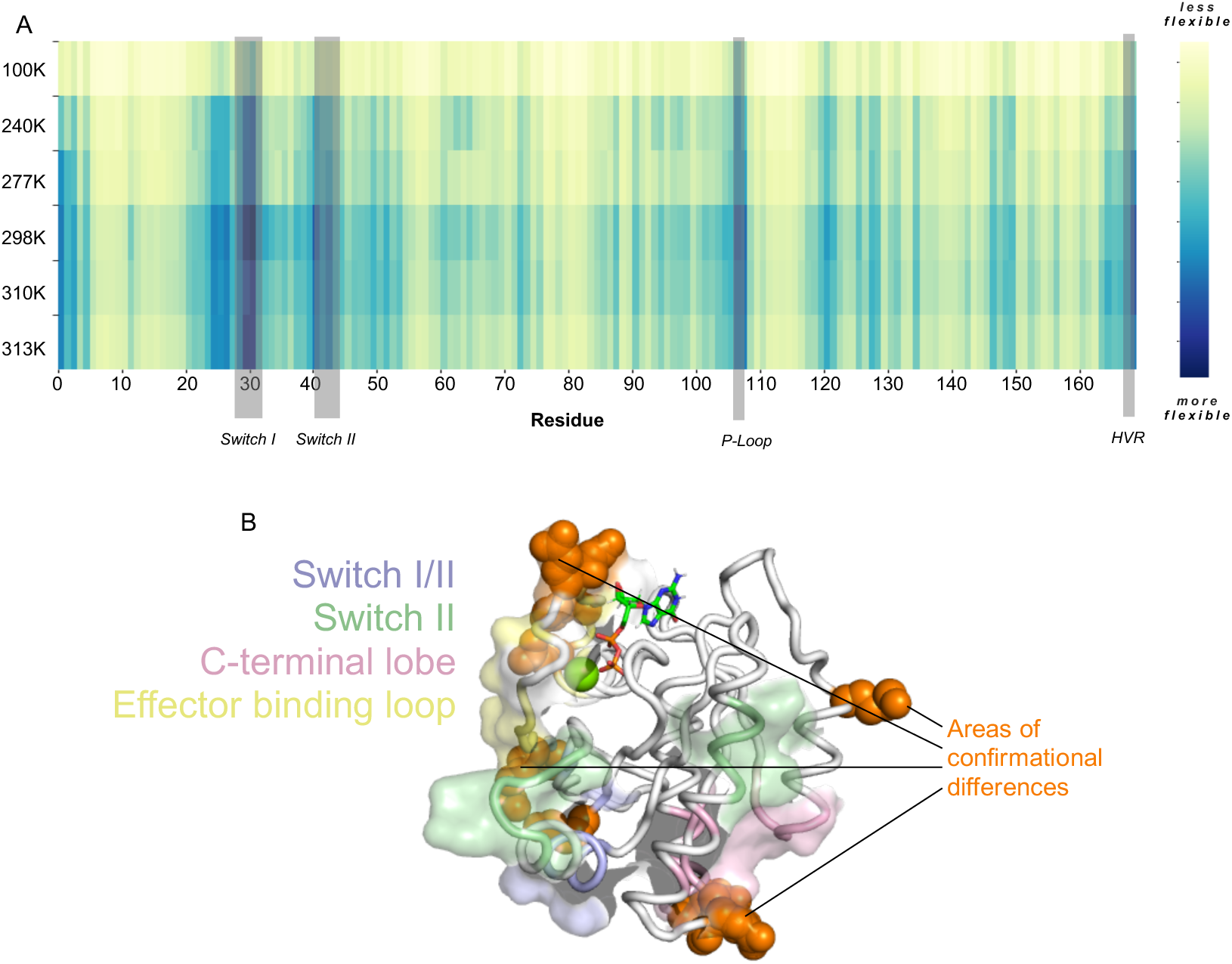
Conformational Analysis and Key Residue Conformations for K-RAS G12C Bound to GDP. ***A***, Root mean square fluctuation heat map (RMSF) of K-Ras G12C bound to GDP across different temperatures plotted per residue. More flexible regions are in blue and key structural regions are labeled and indicated in grey. ***B***, Structure of KRAS G12C at 310K shown with known pocket regions colored. Switch I/II pocket (blue), Switch II pocket (green), C-terminal lobe pocket (pink), and the Effector binding loop pocket (yellow). Residues exhibiting significant conformational differences at 310K (human body temperature) compared to other temperatures are highlighted as orange spheres.

Our analysis identified several key regions that exhibit significant deviations in flexibility as the temperature increases. Residues within the known allosteric and drug pockets showed marked increases in B-factor values at increased temperatures, reflecting a more dynamic structural landscape. Intriguingly, these regions correspond to allosteric and drug-binding sites on K-RAS, as previously predicted from a large-scale mutational screen (35) (Fig. 4B). This finding not only confirms the locations of these critical sites but also provides a structural framework for pinpointing transient allosteric pockets that may serve as alternative therapeutic targets. By mapping these dynamic conformations, our work offers a model for rational drug design that incorporates temperature-dependent structural plasticity. Future studies leveraging high-resolution room-temperature crystallography or molecular dynamics simulations could further refine our understanding of these transient structural elements and their implications for drug development.

### Evaluating Binding Mode Stability of Approved K-RAS G12C Inhibitors

To see whether clinically approved K-RAS G12C inhibitors exhibit temperature-dependent binding variations, we determined the structures of K-RAS G12C at RT in complex with Sotorasib and Adagrasib, the first FDA-approved covalent inhibitors targeting this mutation (37, 38). We observed no significant differences in inhibitor binding poses. In all cases, both Sotorasib and Adagrasib consistently engaged the mutant cysteine residue, locking K-RAS into an inactive conformation. This invariance suggests that these inhibitors represent an optimized binding mode that remains thermodynamically favorable across different conditions, likely due to their covalent mechanism of action. Given that these inhibitors irreversibly engage K-RAS G12C, their high binding affinity and structural rigidity likely limit conformational variability, reinforcing why they remain the dominant therapeutic drugs for targeting this mutant variant.

We then examined whether noncovalent inhibitors designed against other K-RAS mutants might exhibit temperature-dependent binding dynamics. Specifically, we analyzed MRTX-1133, a noncovalent inhibitor targeting K-RAS G12D, which is selective for the inactive GDP-bound state (Fig. 5A) (39). Structural comparisons of cryogenic and RT structures of MRTX-1133 bound to active (GMP-PNP-bound) and inactive (GDP-bound) K-RAS G12D revealed notable flexibility within the C2 moiety of the inhibitor, particularly in the activated conformation (Fig. 5B). This suggests that unlike covalent inhibitors, which irreversibly stabilize K-RAS in a single binding pose, noncovalent inhibitors may exploit dynamic conformational states, potentially leading to varying potency across different conditions. By leveraging this approach, future drug optimization efforts could focus on designing inhibitors that accommodate K-RAS conformational plasticity, maximizing binding efficacy across diverse oncogenic states or targeting either GDP or GTP bound K-RAS.

**Figure 5.**
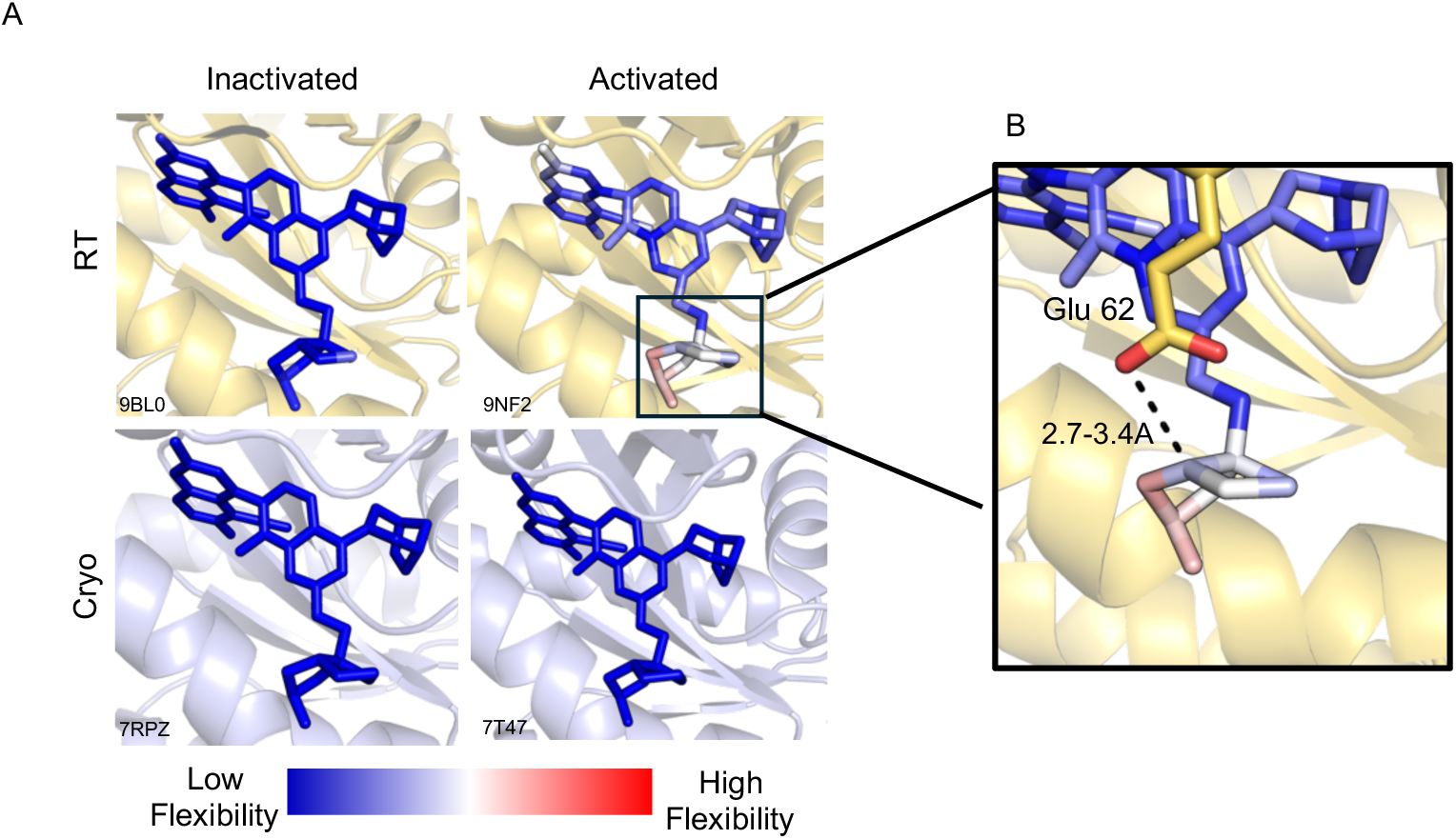
MRTX-1133 exhibits different flexibility when bound to activated versus inactivated K-RAS G12D. ***A,*** MRTX-1133 complexed to either the signaling-active (GMP-PNP) or signaling-inactive (GDP) bound K-RAS G12D under RT or cryogenic temperatures. MRTX-1133 flexibility is shown with blue (low) to red (high) flexibility. ***B,*** A zoomed-in view of the pyrido[4,3-d] pyrimidine substituent at the C2 position of MRTX-1133 showing its flexibility when interacting with Glu 62 (yellow). The H-bonding distance (dashed line) is 2.7-3.4 Å between the C2 component to the residue, as determined by alternative conformations determined with qFit.

## Discussion

In this study, we explored the structural dynamics of WT and mutant K-Ras proteins using multi-temperature X-ray crystallography (MT-XRC), focusing on the implications that temperature-sensitive conformational changes hold for new drug discovery. Mapping allosteric sites through MT-XRC is becoming an increasingly important tool in drug discovery, especially for targets that were once deemed undruggable. This technique makes it possible to capture structural states that emerge exclusively at physiological or elevated temperatures, providing a foundation for targeting proteins that have eluded conventional inhibitor strategies. By incorporating temperature-dependent structural studies into early-stage drug discovery, we can gain deeper insights into the conformational landscapes that drive protein function. Our findings provide new insights into the flexibility and variability of K-RAS, which are necessary for gaining a comprehensive understanding of its function in signaling pathways and for designing new targeted inhibitors.

We determined the high-resolution RT structure of GDP-bound WT K-RAS at 1.4 Å resolution, capturing temperature-dependent structural features in the nucleotide-binding pocket, the Switch I and II domains, and the P-loop. These regions exhibited enhanced flexibility at RT compared to their cryogenic counterparts, suggesting that cryogenic conditions may mask conformational states relevant to K-RAS function. Importantly, we observed that the nucleotide-binding pocket showed subtle shifts in residue positions, particularly in the P-loop (residues 10–17) and Switch I (residues 25–41). These dynamic regions are essential for the ability of K-RAS to transition between active and inactive states, which is critical for signaling.

When comparing the structure of WT K-RAS with that of the oncogenic G12C mutant at RT, we noted significant structural perturbations in several key regions, including residues 1, 4, 23–26, 30, 46–51, 121, 122, 126, and 169. These changes suggest that the G12C mutation induces conformational alterations that could influence nucleotide exchange and effector binding, as well as its known effects on GTP hydrolysis, key processes in K-RAS-driven oncogenesis. Our temperature-dependent studies further revealed that the G12C mutant exhibits increased structural variability at higher temperatures, highlighting the role of temperature in modulating protein dynamics and drug-binding landscapes.

One of the most intriguing consequences of this study is the potential role that temperature-dependent crystallography can play in new drug discovery. By analyzing the structure of K-RAS G12C across a temperature gradient from cryogenic (100K) to physiological (310K) and fever-like (313K) conditions, we observed that structural flexibility increased with temperature. This is particularly evident in regions like the P-loop, Switch I, and Switch II, which are crucial for GTP binding, hydrolysis, and effector interactions. These findings emphasize the importance of studying proteins at physiologically relevant temperatures, as temperature-dependent dynamics can reveal cryptic drug-binding sites that may not be apparent under cryogenic conditions. The temperature-induced conformational changes may help explain why some inhibitors fail or succeed in targeting specific K-RAS oncogenic mutants, highlighting the need to consider protein flexibility in drug design.

For example, using MT-XRC, we identified additional druggable pockets in the G12C mutant that were not visible in the cryogenic structures, including the pocket targeted by the FDA-approved covalent K-RAS G12C inhibitors, Sotorasib and Adagrasib. Intriguingly, these regions correspond to known allosteric and drug-binding sites on K-RAS, as previously identified in a screen of 20,000+ mutations (35). While mutational studies highlight functionally important residues, MT-XRC crystallography offers a powerful complement to this approach by providing precise structural snapshots, capturing the exact conformations of these critical allosteric sites in unprecedented detail. This structural insight enables a deeper understanding of how to target transient allosteric sites, which may play a role in drug resistance or aid in the development of next-generation inhibitors. Given that MT-XRC crystallography reveals a greater number of biologically relevant conformations, it offers a more accurate representation of the protein’s dynamic landscape, essential for designing inhibitors that can adapt to different conformational states of K-RAS.

Although our current study focused on the G12C mutation of K-RAS, other mutations such as G12D, which occur in a wide variety of cancers, will be important to examine in future studies. The addition of G12D data to our structural analysis could provide valuable insights into the broader conformational landscape of K-RAS oncogenic mutants, which will be critical for developing inhibitors that can target multiple K-RAS mutations effectively. We envision future studies where G12D is incorporated into our current framework, either by extending the current analysis or through additional mutagenesis experiments to further assess its impact on K-RAS structure and function.

Our findings underscore the importance of temperature in studying the structure of K-RAS and offers a compelling argument for the integration of MT-XRC crystallography into drug discovery efforts. By revealing conformational flexibility and new druggable pockets that are otherwise obscured at cryogenic temperatures, these studies open up new possibilities for the rational design of inhibitors targeting K-RAS. Thus, in the future we plan to further investigate how temperature-dependent structural changes influence the binding of both covalent and non-covalent inhibitors, as well as explore the potential for targeting alternative allosteric sites in G12C and other K-RAS mutants, and ultimately leverage the dynamic structural insights provided by RT crystallography to examine other GTP-binding proteins that function as molecular switches in cell signaling,.

## Experimental Procedure

### Purification of K-RAS Bound to GDP

Plasmids of both Wild-type and mutant versions of human K-RAS 4B, amino acids 1-169 of human K-RAS 4B, were obtained from Addgene (#111849, #159439, #111850, #111848) which contained a hexa-histidine-tagged recombinant form of human K-RAS (amino acids 1-169) and were transformed into *Escherichia coli* BL21 (DE3) cells. An overnight culture of bacteria was grown at 37°C and used to inoculate 4L of Terrific broth containing 30 mg/ml kanamycin until an OD_600_ of 0.4-0.6 was reached. When the cultures reached an OD_600_ of 0.4-0.6, they were cooled, induced with 0.5 mM IPTG, and grown overnight at 16°C. The cells were then pelleted the next morning and either used immediately or stored at −80°C.

The cell pellet was resuspended in lysis buffer (20 mM Tris pH 8.0, 500 mM NaCl, and 5 mM imidazole) containing protease inhibitors (Leupeptin, Aprotinin, Pepstatin A, AEBSF at 1 mg/mL 1000X stocks). Resuspended cells were lysed with two passes through an Emulsiflex at 15,000 psi, 4°C. After lysis, 2 mM β-mercaptoethanol (βME) was added to the lysate. Debris was removed by ultracentrifugation at 35K, 4°C for 1 hour in a Ti45 rotor. The supernatant was removed and incubated with 5 mL of packed Cobalt Agarose Beads (GoldBio) with end-over-end rotating for an hour before being applied to a column, allowed to settle, and washed 10 CVs with a lysis buffer containing 2 mM βME. The bound protein was then eluted using 5 CVs of elution buffer (20 mM Tris 8.0, 300 mM NaCl, 250 mM imidazole) and dialyzed in 3.5k MWCO Snakeskin dialysis tubing (Thermo Fisher Scientific) overnight in dialysis buffer containing 300 mM NaCl, 20 mM Tris, pH 8.0, 5 mM imidazole, 1 mM dithiothreitol (DTT) and 0.5 mM EDTA. 1 mg GDP per 20 mg K-RAS and 1 mg of hexa-histidine-tagged TEV protease was added before dialysis.

The cleaved protein was then applied to a 5 mL HisTrap™ FF Column (Cytiva) to remove any TEV protease and un-cleaved protein. The protein solution was diluted five-fold into dilution buffer containing 50 mM NaCl, 20 mM Tris, pH 8.0 before applying it to an anion exchange column (2X Q HiTrap FF 5 mL connected in tandem) with a 50-500 mM salt gradient (including 20 mM Tris, pH 8.0). The protein containing peaks were combined and concentrated before being applied to a Superdex 75 10/300 GL size exclusion chromatography column equilibrated in 20 mM HEPES, pH 7.5, 150 mM NaCl and 1 mM DTT buffer.

### Purification of K-RAS G12D bound to GMPPNP

The same procedure for K-RAS G12D bound to GDP was followed until after the Q column chromatography was completed (GMPPNP was used instead of GDP before dialysis). At this point, 1.5 mL of ion exchange-purified RAS protein was mixed with 3 mg of GMPPNP (~4 mM final). EDTA was added to a final concentration of 25-30 mM. After incubation for 1 hour at RT, the reaction buffer was exchanged for phosphatase buffer (32 mM Tris, 20 mM Ammonium Sulfate, 0.1 mM ZnCl2 pH = 8.0) by dialysis using a 10kDa MWCO Slide-A-Lyzer dialysis cassette (MWCO 10,000 Da). Thirty units of Calf Intestinal Phosphatase (CIP, NEB) was then added with 2 mg of GMPPNP. After a 1-hour incubation at 22 °C, MgCl_2_ was added to the final concentration of 30 mM. The mixture was incubated on ice for 15 min before being concentrated using a Amicon-4 concentration (10,000 MWCO) to ~700 µL. Finally, the solution was applied to a Superdex 75 10/300 GL for size exclusion chromatography (buffer includes 20 mM HEPES 7.5, 150 mM NaCl, 1 mM MgCl_2_) and the purified protein was concentrated before flash freezing and being stored at −80°C. HPLC analysis was done to verify the nucleotide status as previously described (Fig. S2) (46).

### Crystallography of K-RAS bound to GDP and GMPPNP

For crystallization of K-RAS bound to GDP and MRTX-1133, following gel filtration, the purified protein was concentrated to 20, 30, and 40 mg/mL before MRTX-1133 was added in 4-fold molar excess. The protein and drug were then centrifuged at high speed to remove any aggregates. Crystals were grown using hanging drop vapor diffusion in 24 well trays in mother liquor containing 0.1 M Bis Tris pH=5.5, 0.1 M NaOAc pH 3.0-5.6, 8% v/v 2-propanol, and 24% PEG 4000. Similar conditions were used to obtain crystals of K-RAS bound to GDP and Sotorasib or Adagrasib (Medchemexpress).

For K-RAS bound to MRTX-1133 and GMPPNP, following gel filtration, the purified protein was concentrated to ~18 mg/ml before MRTX-1133 was added in 3-fold molar excess. The samples were then centrifuged at high speed to remove aggregates. Crystals were also grown using hanging drop vapor diffusion in 24 well trays in mother liquor containing 25% PEG, 100 mM ammonium acetate, 0.1 M sodium citrate (pH=4.2-6.0).

The crystals were then analyzed at the Cornell High Energy Synchrotron Source (CHESS) HPBio MX & FlexX beamline at cryogenic and room temperature. The data was processed using HKL2000 (40) and Phenix (41) and phased against PDB ID: 7RPZ (39). These structures were then visualized using PyMOL (42) and validated using MolProbity (43) and wwPDB validation software before being deposited into the PDB. The flexibility of MRTX-1133 was examined using qFIT (44).

## Data, Materials, and Software Availability

Atomic coordinates and structure factors for the reported K-RAS structures have been deposited in the Protein Data Bank (PDB). All other relevant data are available upon request from the corresponding author. Software used in the analysis includes PyMOL, CCP4 (45), PHENIX, which are available through their respective distribution channels. Experimental materials, including mutant plasmids, are available upon request for non-commercial research purposes.

## Author Contributions

S.D. conceived and designed the experiments, with S.D. and M.X. conducting the experimental work. Sample preparation was a collaborative effort between S.D., M.X., and S.M. All authors contributed to interpreting the results. S.D. drafted the manuscript, with valuable input from all authors. Additionally, each author provided critical feedback and helped refine the research, analysis, and manuscript.

## Acknowledgements

This work was supported by NIH grant to RAC EY034867 from the National Eye Institute. SD acknowledges support from NSF GFRP and the Sloan Foundation. X-ray crystallography was conducted at the Center for High-Energy X-ray Sciences (CHEXS), supported by the National Science Foundation (BIO, ENG and MPS Directorates) under award DMR-1829070, and the Macromolecular Diffraction at CHESS (MacCHESS) facility, by 1-P30-GM124166-01A1 from the National Institute of General Medical Sciences, National Institutes of Health, and by New York State’s Empire State Development Corporation (NYSTAR). This research also used resources of the National Synchrotron Light Source II, a U.S. Department of Energy (DOE) Office of Science User Facility operated for the DOE Office of Science by Brookhaven National Laboratory under Contract No. DE-SC0012704. We would also like to thank Kevan Shokat for providing us with MRTX-1133.

---

**Table 1.**
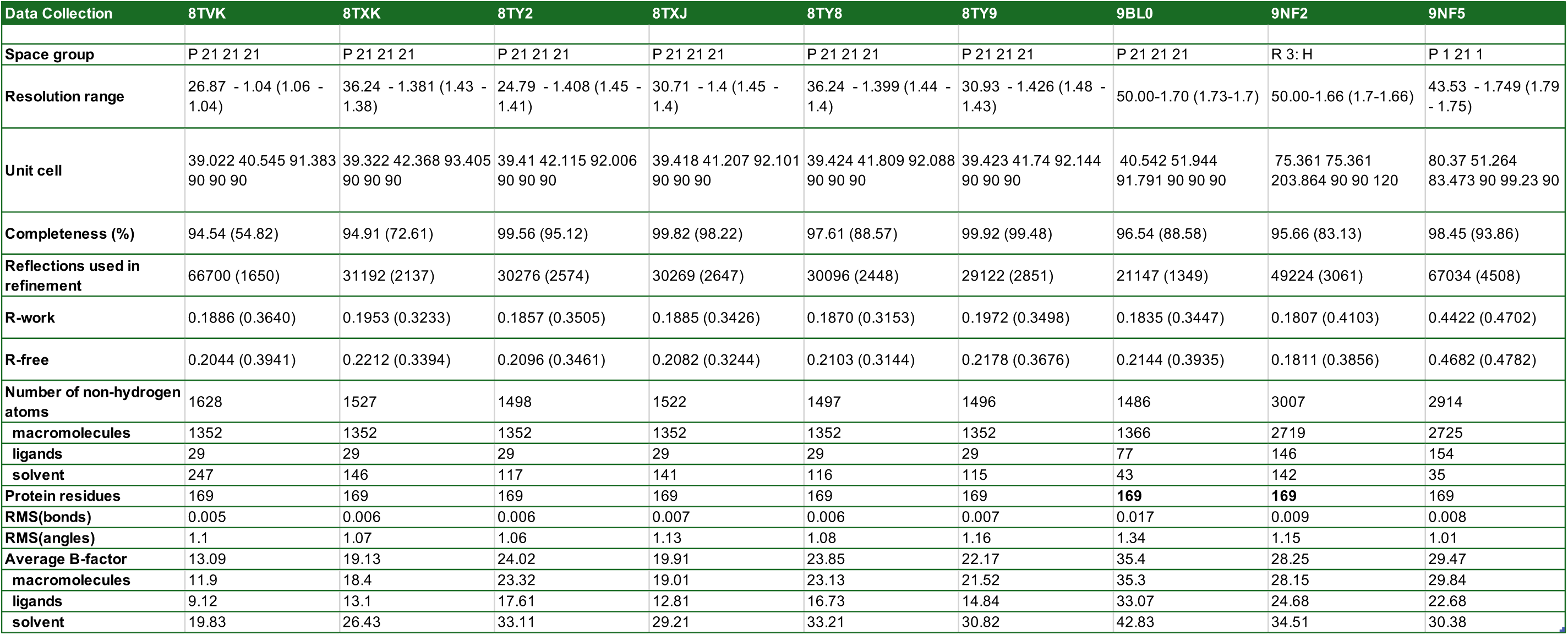
Crystallization and Data Collection Statistics for Solved K-RAS Structures.

