## Supplementary Figures for "Revealing Functional Hotspots: Temperature-Dependent Crystallography of K-RAS Highlights Allosteric and Druggable Sites"

### Supplementary Figures and Figure Legends

#### Supplementary Figure 1

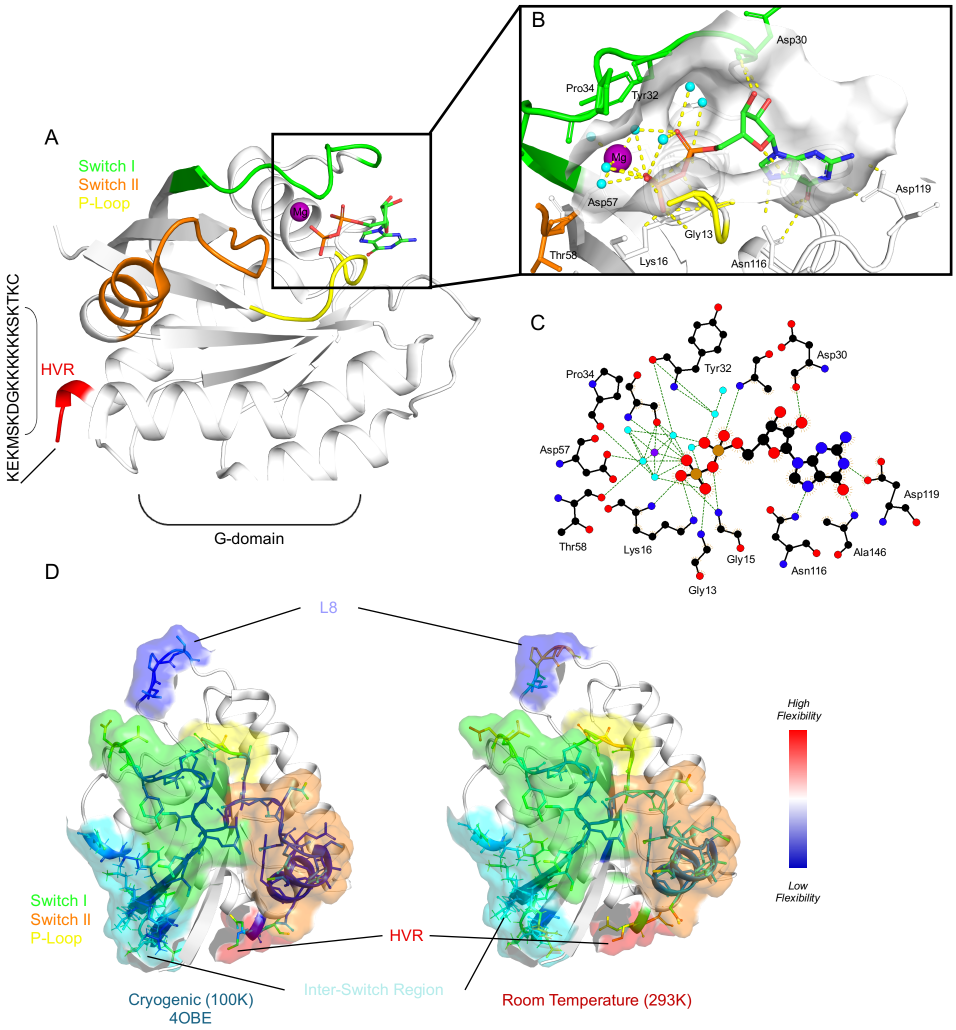

**Supplementary Figure 1. The Room Temperature (RT) Crystal Structure of K-RAS at 1.4 Å.** ***A***, The G-domain (amino acids 1-169) of WT K-RAS bound to GDP at RT in cartoon representation with Switch I (green) and II (orange), P-loop (yellow), and Hyper Variable Regions (HVR) (red) highlighted, showing clear structural definition across all regions. ***B***, Zoomed-in region of the nucleotide-binding pocket with GDP. Magnesium (purple sphere), coordinating waters (blue spheres), the Switch regions, and the P-loop are shown. ***C***, Schematic representation of the interactions between GDP and WT K-RAS, generated using LigPlot+. The figure highlights key hydrogen bonds (represented as dashed lines) stabilizing the GDP molecule within the nucleotide-binding pocket of K-RAS. The magnesium ion (purple sphere) coordinating the phosphate groups of GDP is also shown. Key residues from K-RAS involved in these interactions are labeled. ***D***, Comparison of the RT, 1.4 Å structure of WT K-RAS with its cryogenic (100K) counterpart. Switch I (green) and II (orange), and P-loop (yellow), (HVR) (red), Inter-Switch (blue), and Loop 8 (dark blue) regions are highlighted. Residues are shown, colored from Blue (lowest) to Red (highest) B-factor values, indicating how flexible the regions are.

Supplementary Figure 2

A

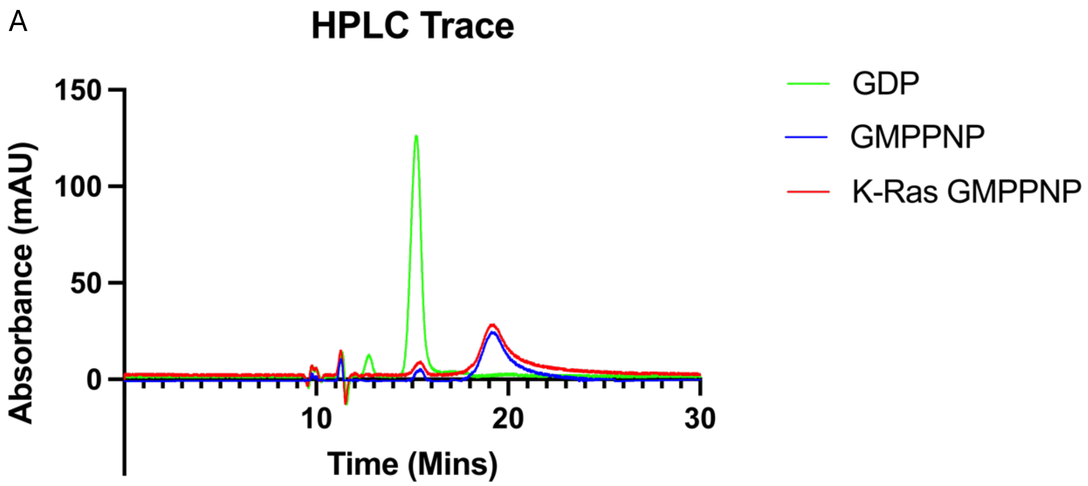

**Supplementary Figure 2. HPLC Chromatogram Showing K-Ras G12D Exchanged with GMPPNP.** ***A,*** Displays a representative chromatogram that monitors UV absorbance at 252 nm, highlighting the nucleotide standards GDP, GMPPNP, and the exchanged K-Ras G12D. Well-resolved peaks are observed for GDP (green line), GMPPNP (blue line), and the exchanged K-Ras (red line).

Supplementary Figure 3

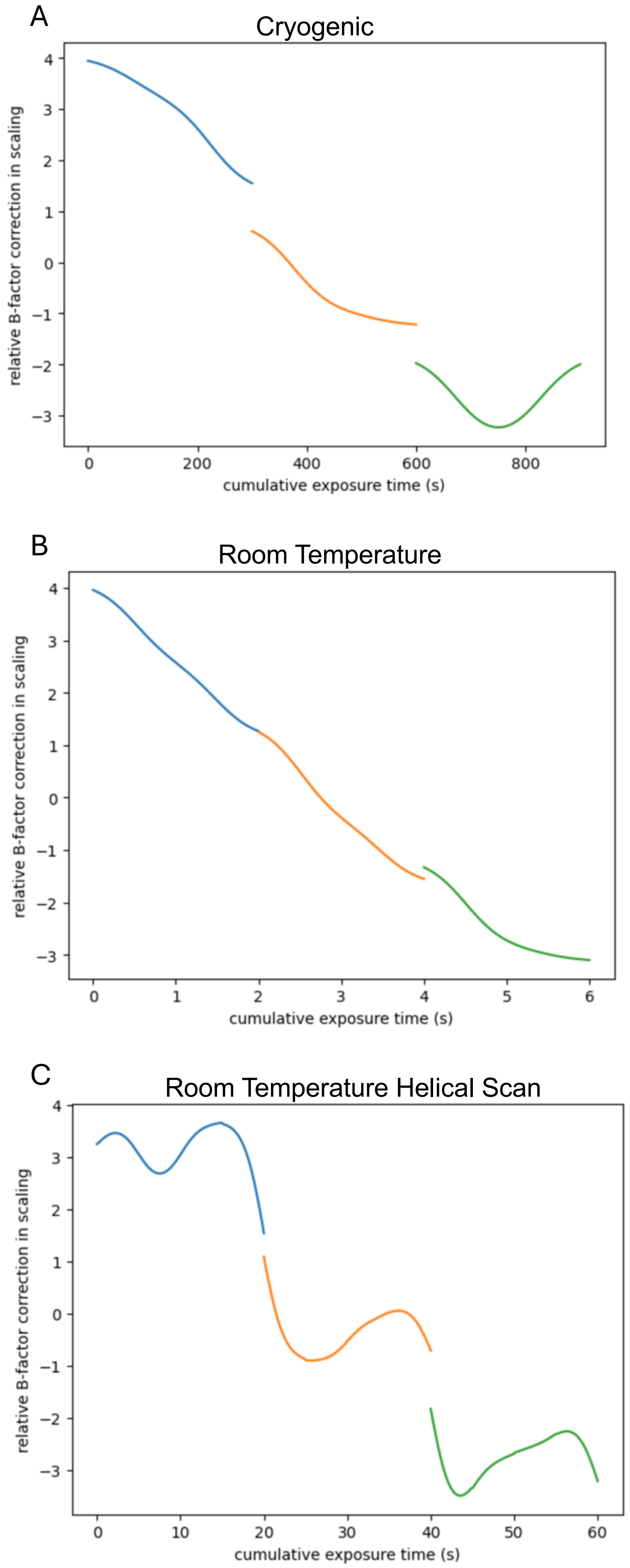

**Supplementary Figure 3. B-Factor Decay Plots of Radiation Damage in K-Ras Protein Structures** ***A,*** B-factor decay at cryogenic temperatures (100K), exhibiting minimal radiation damage. Blue, Orange, and Green line represent consecutive data collections over 10 degree at the same crystal spot. ***B,*** B-factor decay at room temperature (298 K) ***C,*** B-factor decay at room temperature (298K with helical scanning.
